## Supplemental Figures 1-12 for "SCC3 is an axial element essential for homologous chromosome pairing and synapsis"

**Supplemental Information**

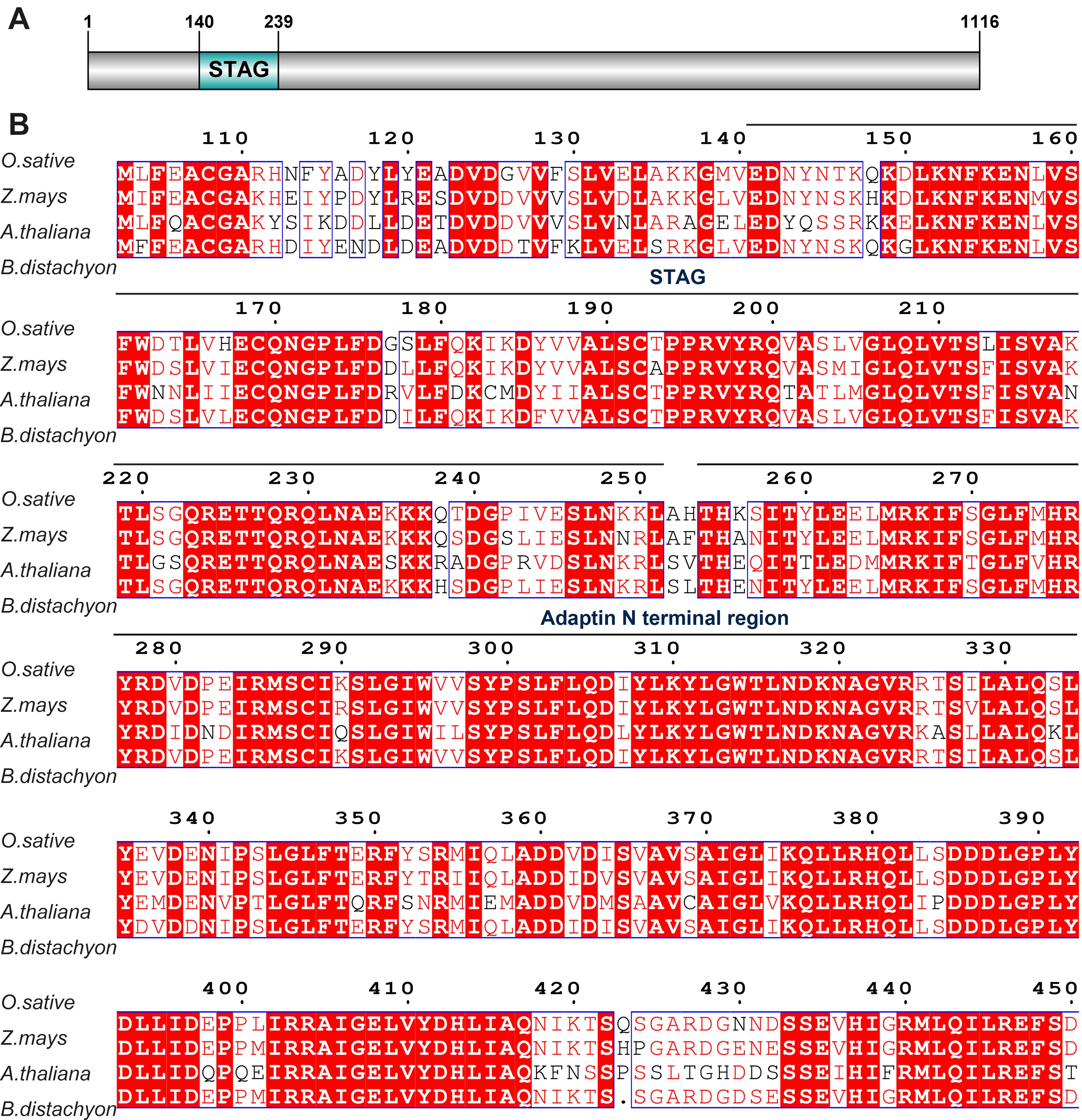

**Figure S1 (related to Figure 1). Multiple sequence alignment of SCC3 with its homolo****gs from three different species**

(A) SCC3 contains a conserved STAG domain.

(B) STAG domains are conserved in both monocotyledonous and dicotyledonous plants.

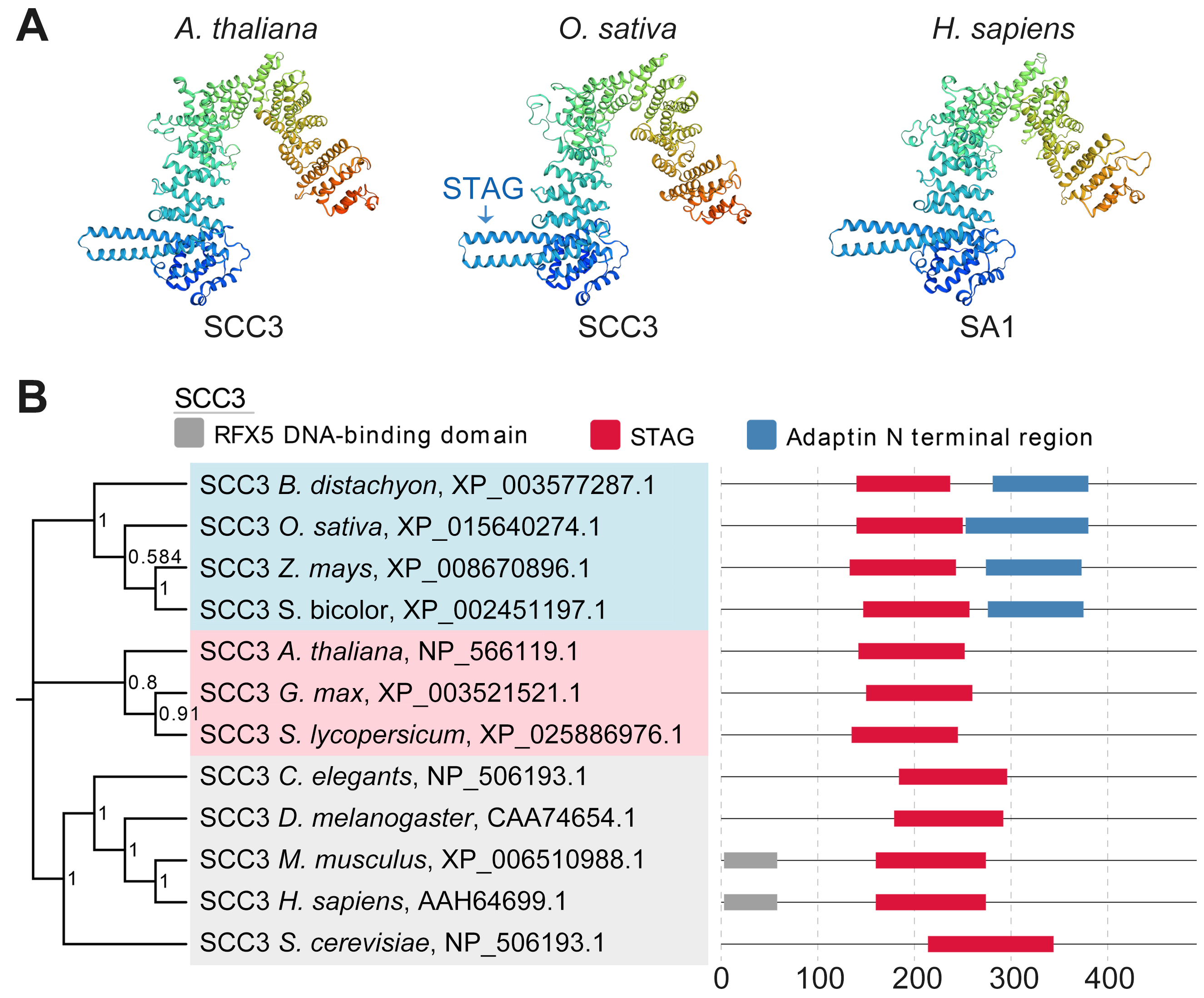

**Figure S2 (related to Figure 1). Protein structure and phylogenetic tree analysis of SCC3**

(A) The full-length (1–1116 aa) SCC3 protein among different species shows high structural similarity. The arrow indicates similarity in *SCC3* subunits.

(B) Phylogenetic tree derived from full-length SCC3 amino acid sequences and homologous sequences from other plant species. The right panel indicates the conserved domains of SCC3 proteins in different species.

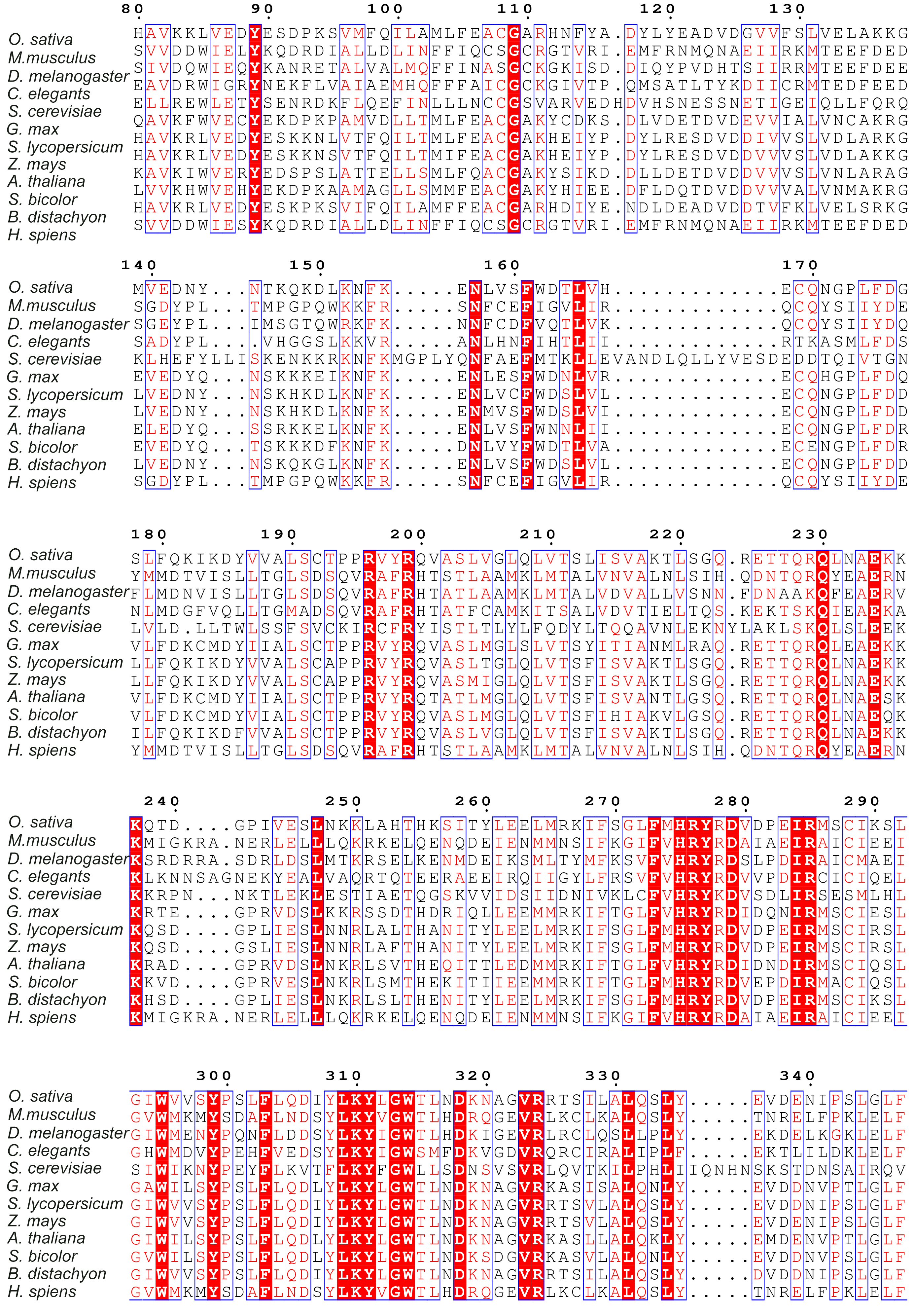

**Figure S3 (related to Figure 1). Multiple sequence alignment of SCC3 with its homologs from different species analyzed in evolutionary tree**

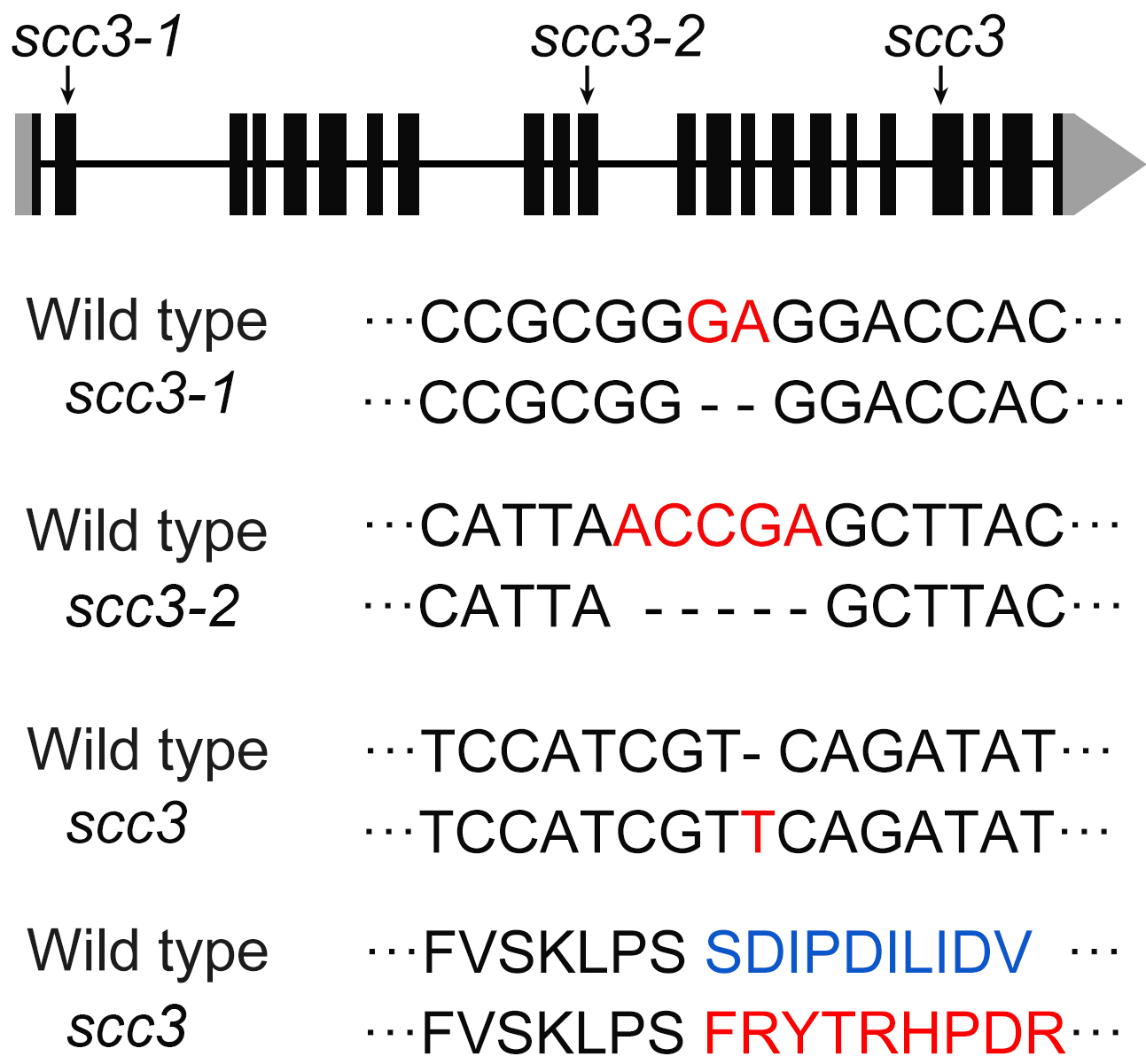

**Figure S4 (related to Figure 1). Schematic representation of SCC3’s mutation sites**

Schematic representation of the *SCC3* gene and its mutation site. Coding regions are shown as black boxes and untranslated regions are shown as gray boxes.

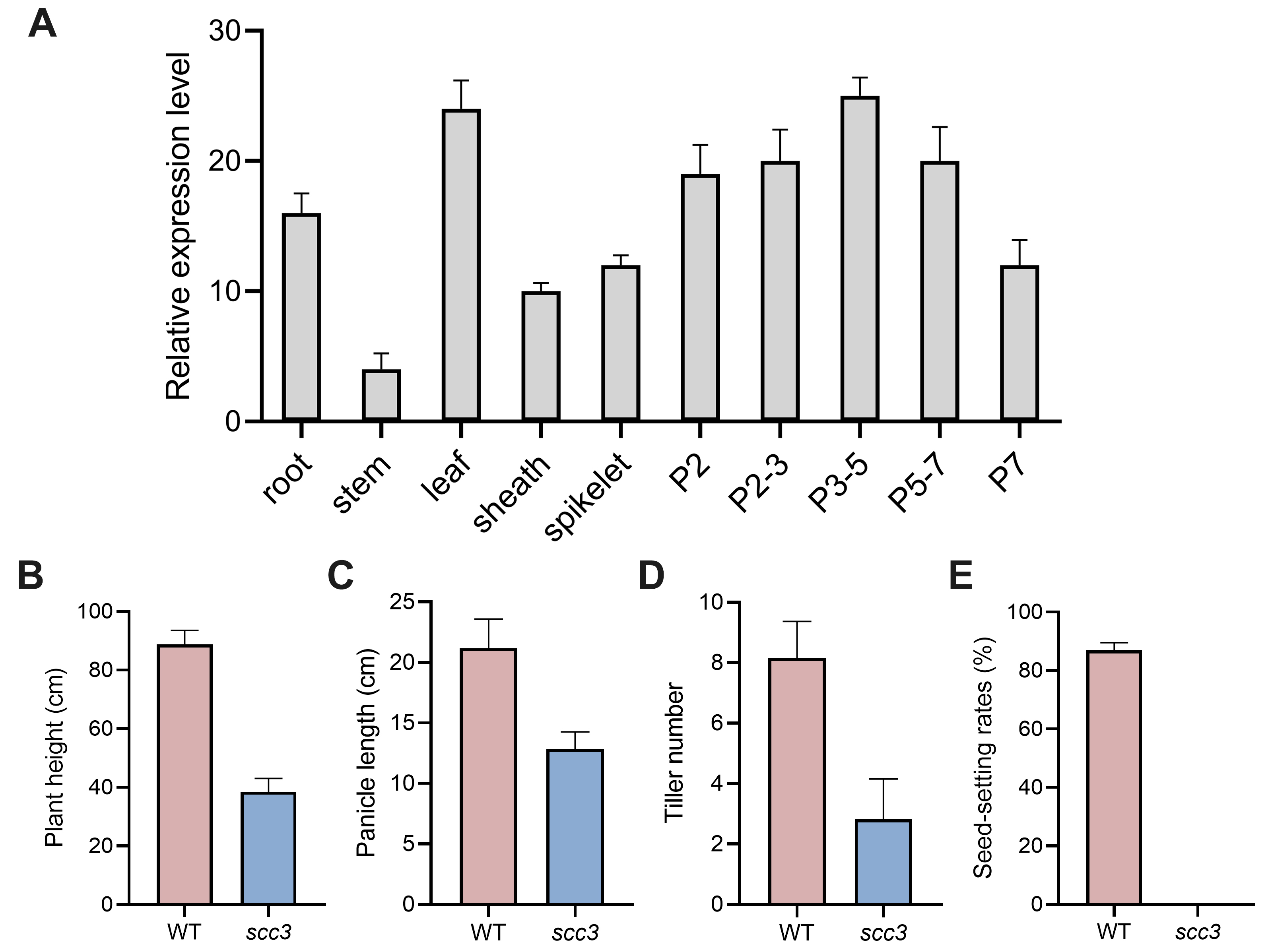

**Figure S5 (related to Figure 1). The expression pattern of SCC3 and plant phenotypic statistics of wild type and *scc3* mutants**

(A) The expression pattern of *SCC3*. P2, 2 cm long panicles; P2-3, 2 to 3 cm long panicles; P3-5, 3 to 5 cm long panicles; P5-7, 5 to 7 cm long panicles; P7, 7 cm long panicles. Expression values represent means ± SD of three biological replicates.

(B) Statistics of plant height in wild type and *scc3*.

(C) Statistics of panicle length in wild type and *scc3*.

(D) Statistics of tiller number in wild type and *scc3*.

(E) Statistics of seed-setting rates in wild type and *scc3*.

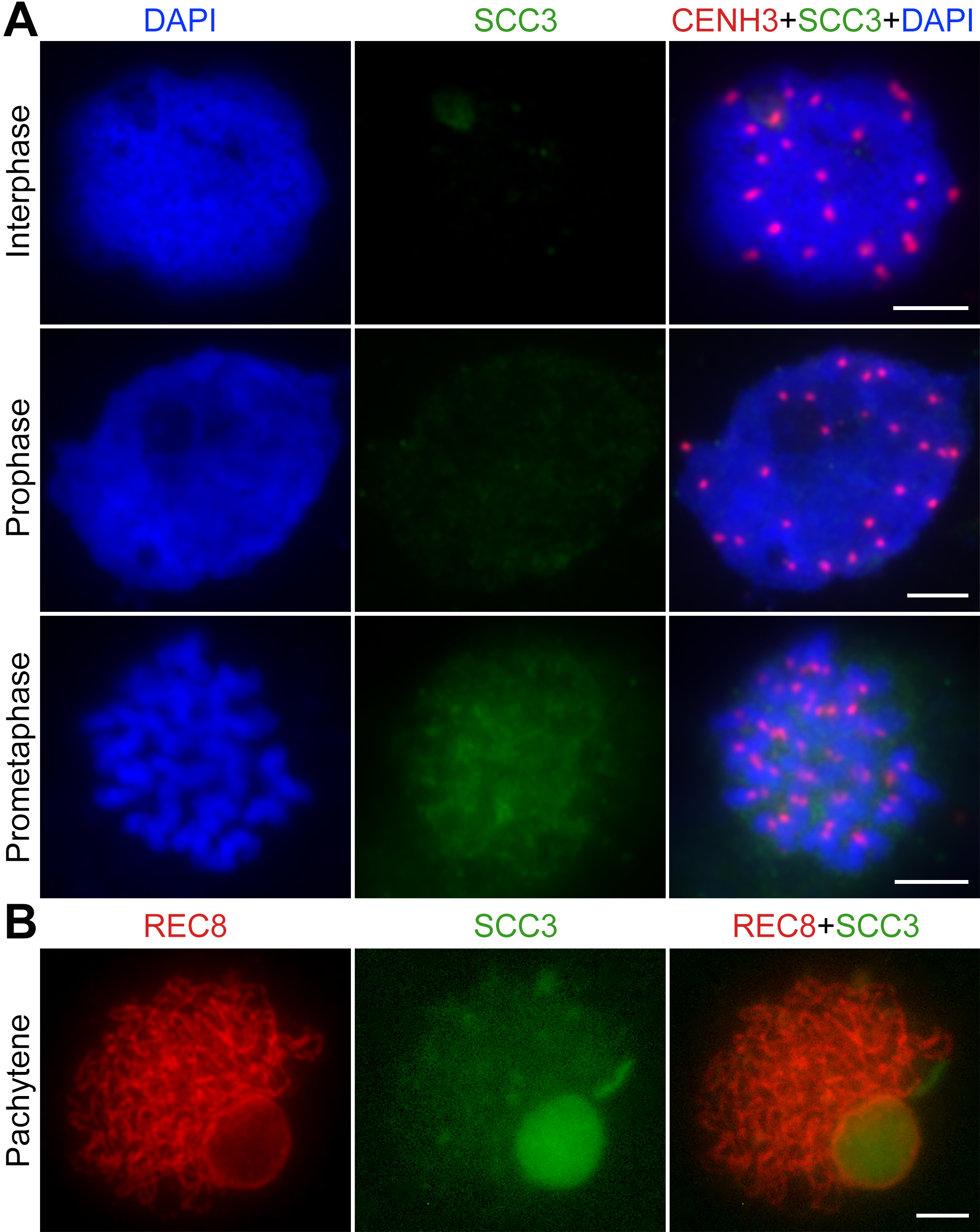

**Figure S6 (related to Figures 2 and 3). Immunolocalization of SCC3 and REC8 in *scc3* mitosis and meiosis**

(A) Immunolocalization of CENH3 (red, from rabbit) and SCC3 (green, from mouse) in *scc3* mutant at mitosis. Chromosomes were stained with DAPI. Bar, 5 μm.

(B) Immunolocalization of REC8 (red, from rabbit) and SCC3 (green, from mouse) in *scc3* meiocytes during meiosis. Bar, 5 μm.

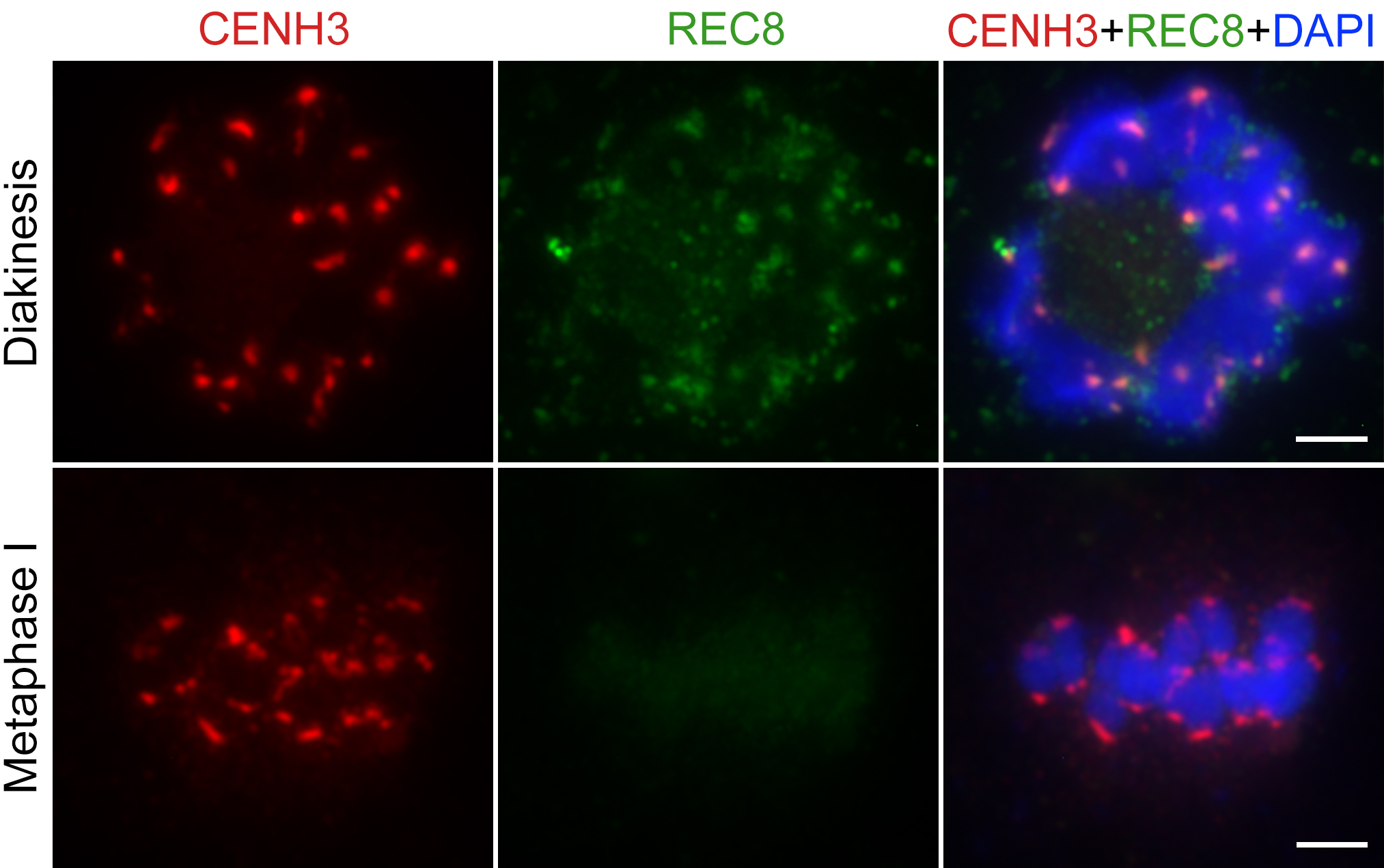

**Figure S7 (related to Figure 3). Immunolocalization of CENH3 and REC8 in wild type from diakinesis to metaphase I**

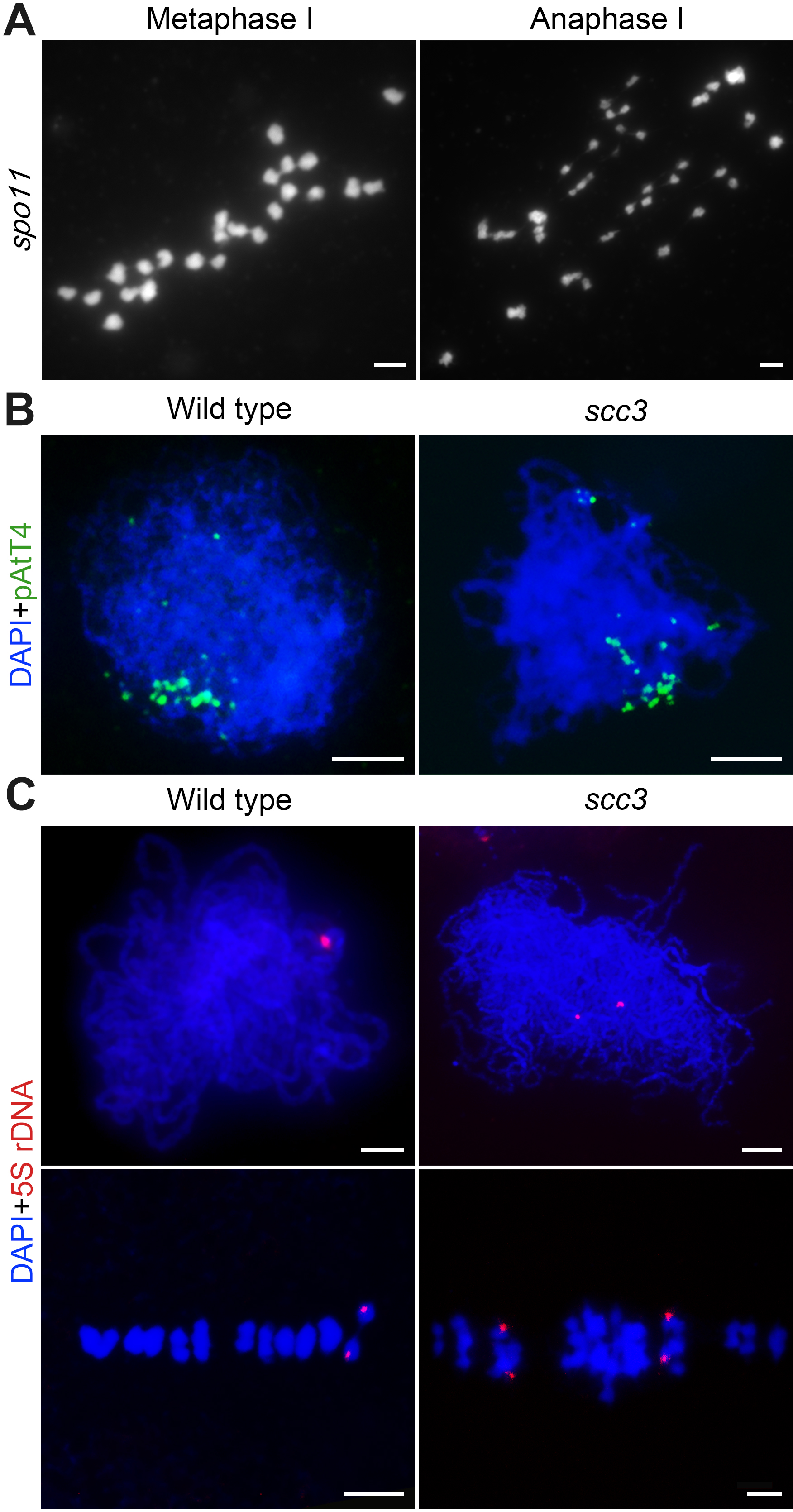

**Figure S8 (related to Figure 5). Chromosome behaviors of *spo11* and cytogenetic analysis of homologous pairing in wild type and *scc3***

(A) Chromosome behavior of *spo11* in metaphase I and anaphase I. Bars, 5 μm.

(B) Telomere bouquets were detected in both wild type and *scc3* with the telomere-specific probe (pAtT4, green) by fluorescence in *situ* hybridization (FISH) assays. Chromosomes were stained with DAPI. Bars, 5 μm.

(C) The pairing status of chromosomes revealed by 5S rDNA in wild type. Chromosomes were stained with DAPI. Bars, 5 μm.
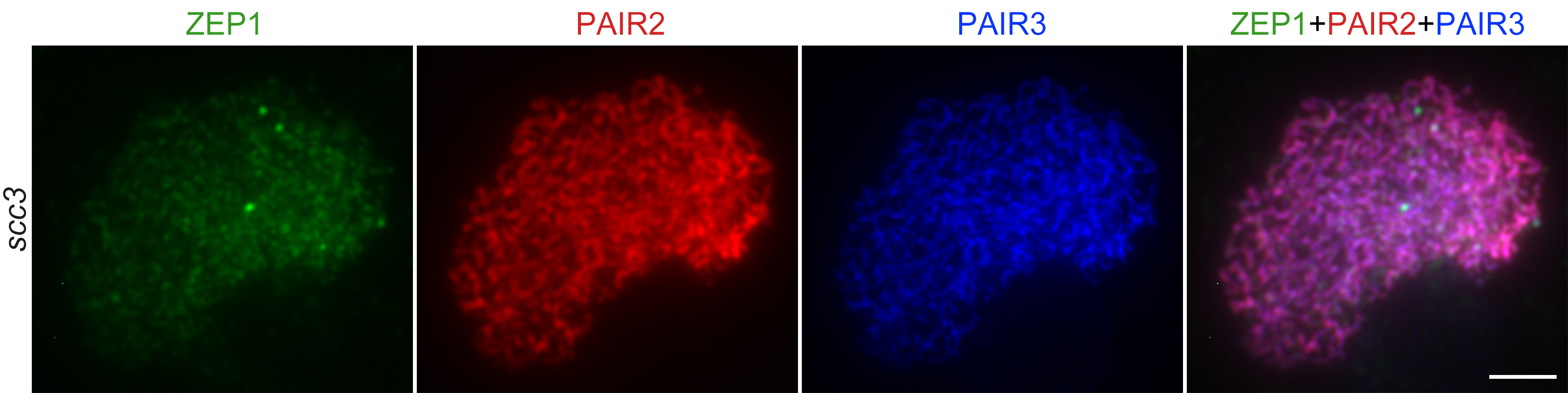

**Figure S9 (related to Figure 5). Immunolocalization of ZEP1, PAIR2 and PAIR3 in *scc3* at zygotene**

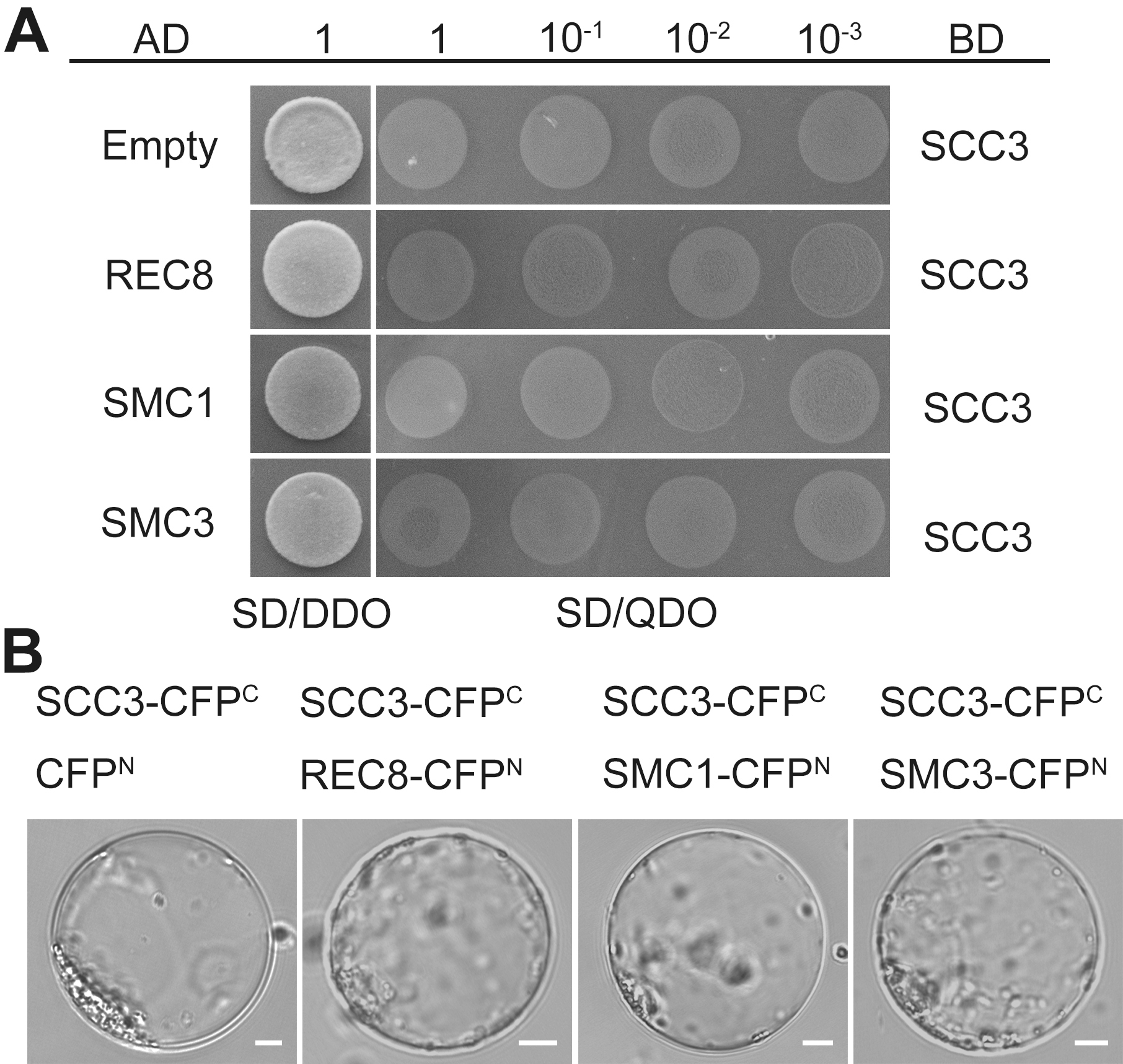

**Figure S10 (related to Figure 7). SCC3 does not interact with other cohesin proteins**

(A) SCC3 does not interact with REC8, SMC1 and SMC3 in yeast-two-hybrid assays.
(B) SCC3 does not interact with REC8, SMC1 and SMC3 in bimolecular fluorescence complementation assays in rice protoplast. Bars, 5 μm.

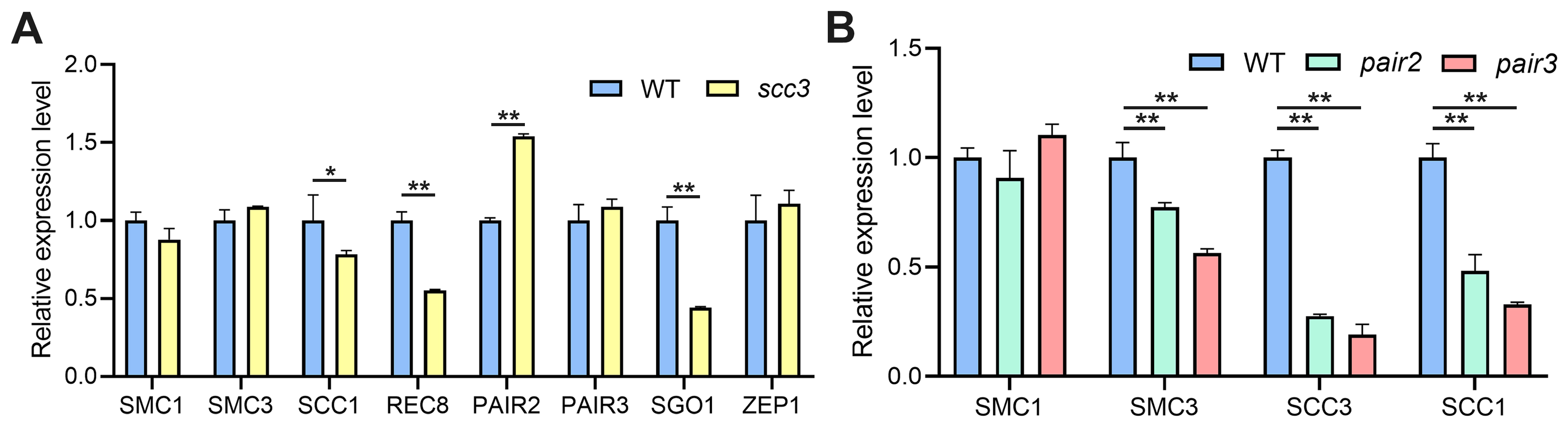

**Figure S11 (related to Figure 7). The expression levels of cohesin-related genes in wild type and other mutants**

(A) The expression levels of SMC1, SMC3, SCC1, REC8, PAIR2, PAIR3, SGO1 and ZEP1 in *scc3* and wild type.

(B) The expression levels of SMC1, SMC3, SCC3 and SCC1 in *pair2*, *pair3* and wild type.

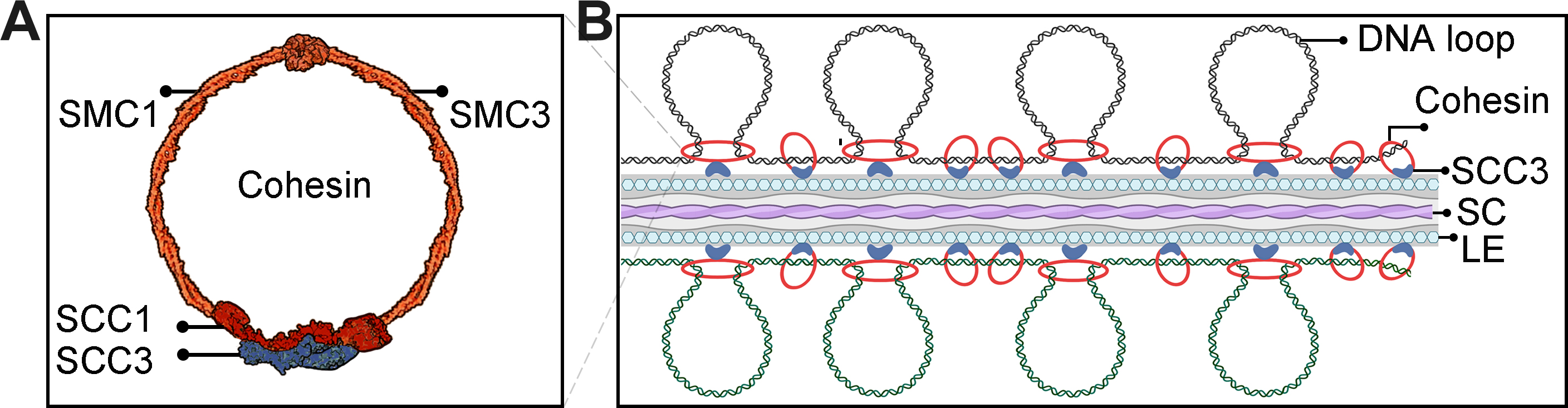

**Figure S12. SCC3 acts as the cohesin and promotes homologous pairing and synapsis**

(A) Structure model of the cohesin complex. The SCC3 subunit is associated with the middle region of SCC1.

(B) During meiosis, SCC3’s localization pattern probably changes to the root of DNA loop with axial elements (AEs), which wraps synaptonemal complex (SC). This indicates SCC3 is a meiotic AE essential for homologous chromosome pairing and synapsis, also affects recombination progress and CO formation.

**Table. S1** Primers for Real-time PCR and plasmid construction

| Primer name | | Primer sequence | Description |
| --- | --- | --- | --- |
| SCC3-Cas9-F | GGCAATTGCGCGAGGCACCAACAT | | CRISPR-Cas9 |
| SCC3-Cas9-R | AAACATGTTGGTGCCTCGCGCAAT | |  |
| REC8-Cas9-F | GGCAGAGCCTTCCGCGCGAGGAGC | | CRISPR-Cas9 |
| REC8-Cas9-R | AAACGCTCCTCGCGCGGAAGGCTC | |  |
| SMC1-RT-F | TAAGGGGACGCAGACAATCG | | Real-time PCR |
| SMC1-RT-R | GGGCGTAGATGAGGTCCTTG | |  |
| SMC3-RT-F | CGTTGTGATCCAGCCCCATT | | Real-time PCR |
| SMC3-RT-R | CTGGTCGGAATGTCGTTGCT | |  |
| SCC1-RT-F | ATGTTCTGTCCGCAATGGGT | | Real-time PCR |
| SCC1-RT-R | TGTCAACACCCTGAATCGCA | |  |
| SCC3-RT-F | GTTATTCATGCACCGCTATCG | | Real-time PCR |
| SCC3-RT-R | CAGGCTTTGCAAGGCAAGAA | |  |
| REC8-RT-F | TCCACTCGTACCTCAAGCTA | | Real-time PCR |
| REC8-RT-R | GTTGCTAAAACGCATGCTTG | |  |
| PAIR2-RT-F | GATCGACGATGGGTTGCCT | | Real-time PCR |
| PAIR2-RT-R | AACAAACGGACGCCTGCTC | |  |
| PAIR3-RT-F | CAAGAAGAGAACAGGACAGT | | Real-time PCR |
| PAIR3-RT-R | GTAGAACGATGAGTAGACAATG | |  |
| SGO1-RT-F | TCAATCAGCTGTGCCATCTT | | Real-time PCR |
| SGO1-RT-R | CATCTTGCCACCACA AATCA | |  |
| ZEP1-RT-F | CAGCAGGATAATGAGCATAA | | Real-time PCR |
| ZEP1-RT-R | GGTCTCAGGACTAACCAACT | |  |
| SCC3-AD-F | GACGTACCAGATTACGCTCATATGGACGAGACCCTAGCCTCC | | Constructing vectors for Y2H experiments |
| SCC3-AD-R | TCTGCAGCTCGAGCTCGATGTCAGCTATTGCTTCCTGATGCCCT | |  |
| SCC3-BD-F | ATCTCAGAGGAGGACCTGCATATGGACGAGACCCTAGCCTCC | | Constructing vectors for Y2H experiments |
| SCC3-BD-R | GCGGCCGCTGCAGGTCGACGTCAGCTATTGCTTCCTGATGCCCT | |  |
| SCC1-AD-F | GACGTACCAGATTACGCTCATATGTTCTACTCGCAGTTCATC | | Constructing vectors for Y2H experiments |
| SCC1-AD-R | TCTGCAGCTCGAGCTCGATGTCAGAAATCTGACTTCAGGAGCTT | |  |
| SCC1-BD-F | ATCTCAGAGGAGGACCTGCATATGTTCTACTCGCAGTTCATC | | Constructing vectors for Y2H experiments |
| SCC1-BD-R | GCGGCCGCTGCAGGTCGACGTCAGAAATCTGACTTCAGGAGCTT | |  |
| SMC1-AD-F | GACGTACCAGATTACGCTCATATGGCCGCGGCGGCGGCAGG | | Constructing vectors for Y2H experiments |
| SMC1-AD-R | TCTGCAGCTCGAGCTCGATGTCACAAAAGAGGTACTTTAA | |  |
| SMC1-BD-F | ATCTCAGAGGAGGACCTGCATATGGCCGCGGCGGCGGCAGG | | Constructing vectors for Y2H experiments |
| SMC1-BD-R | GCGGCCGCTGCAGGTCGACGTCACAAAAGAGGTACTTTAA | |  |
| SMC3-AD-F | GACGTACCAGATTACGCTCATATGGATGCTGAGCGTGACCA | | Constructing vectors for Y2H experiments |
| SMC3-AD-R | TCTGCAGCTCGAGCTCGATGTCAGCTAGCGTTGTGTGTCT | |  |
| SMC3-BD-F | ATCTCAGAGGAGGACCTGCATATGGATGCTGAGCGTGACCA | | Constructing vectors for Y2H experiments |
| SMC3-BD-R | GCGGCCGCTGCAGGTCGACGTCAGCTAGCGTTGTGTGTCT | |  |
| REC8-BD-F | ATCTCAGAGGAGGACCTGCATATGTTCTACTCGCACCAGC | | Constructing vectors for Y2H experiments |
| REC8-BD-R | GCGGCCGCTGCAGGTCGACGCATCTTTGGTCCCCTCGAGAT | |  |
| SCC3-NE/CE-F | CCCAGGCCTACTAGTGGATCCATGGACGAGACCCTAGCCTCC | | Constructing vectors for BiFC experiments |
| SCC3-NE/CE-R | CCCGGGAGCGGTACCCTCGAGTCAGCTATTGCTTCCTGATGCCCT | |  |
| REC8-NE/CE-CF | CCCAGGCCTACTAGTGGATCCATGTTCTACTCGCACCAGC | | Constructing vectors for BiFC experiments |
| REC8-NE/CE-R | ATCCCGGGAGCGGTACCCTCCATCTTTGGTCCCCTCGAGAT | |  |
| SCC3-Ab-F | GACGACGACGACAAGGCCATGCATGTAAGTGATGGGGAAAA | | Antibody production |
| SCC3-Ab-R | GCAGCCGGATCTCAGTGGTGGGCTATTGCTTCCTGATGCCC | |  |
